# Identification and characterization of novel Bovine Leukemia Virus (BLV) antisense transcripts reveals their constitutive expression in leukemic and pre-leukemic clones

**DOI:** 10.1101/039255

**Authors:** Keith Durkin, Nicolas Rosewick, Maria Artesi, Vincent Hahaut, Philip Griebel, Natasa Arsic, Arsène Burny, Michel Georges, Anne Van den Broeke

## Abstract

Bovine Leukemia Virus (BLV) is a deltaretrovirus closely related to the Human T-cell leukemia virus-1 (HTLV-1). Cattle are the natural host of BLV where it integrates into B-cells and produces a lifelong infection. Most infected animals remain asymptomatic but following a protracted latency period about ∽5% develop an aggressive leukemia/lymphoma, mirroring the disease trajectory of HTLV-1. The 5’LTRs of both the BLV and HTLV-1 proviruses are transcriptionally silent in tumors, however they are not entirely quiescent, with the HLTV-1 antisense transcript *HBZ* and the BLV microRNAs constitutively expressed in tumors. Here, using RNA-seq, we demonstrate that in addition to microRNAs, the BLV provirus also constitutively expresses two antisense transcripts in all BLV infected samples examined. The first transcript (AS1) has alternate potential polyadenylation sites generating a short transcript of ∽600bp (aS1-S) and a less abundant longer transcript of ∽2200bp (AS1-L). Alternative splicing also creates a second transcript of ∽400bp (AS2) utilizing the first exon of AS1. Production of AS transcripts from the 3’LTR was supported by reporter assays demonstrating that the BLV LTR has substantial and Tax-independent antisense promoter activity. BLV AS transcripts predominantly localize in the nucleus. Examination of protein coding potential showed AS2 to be non-coding, while the AS1-S/L transcripts coding potential is ambiguous, with a small potential open reading frame (ORF) of 264bp present. The AS1-L transcript overlaps the BLV microRNAs transcribed in the sense direction. Using high throughput sequencing of RNA-ligase-mediated (RLM) 5’ RACE products, we show that the perfect complementary between the transcripts leads to RNA-induced silencing complex (RISC) mediated cleavage of AS1-L. Furthermore, experiments using BLV proviruses where the microRNAs were removed or inverted point to additional transcriptional interactions between the two viral RNA species. Knock down of AS1-S/L using locked nucleic acids (LNAs) showed no obvious effect on the cells phenotype. While a detailed elucidation of the BLV antisense transcripts function remains in the future, the constitutive expression in all samples examined, points to a vital role for the transcripts in the life cycle and oncogenic potential of BLV.

## Introduction

The deltaretrovirus Bovine Leukemia Virus (BLV), like its close relative Human T-cell leukemia virus (HTLV-1), produces a chronic infection in its natural host (cattle and water buffalo), with most infected individuals remaining asymptomatic and about ∽5% going on to develop leukemia/lymphoma [1,2]. In humans the time between infection and the onset of Adult T cell leukemia/lymphoma (ATL) generally spans several decades [3], while in cattle several years separate infection from progression to B-cell leukemia/lymphoma [4]. BLV infections are common in cattle throughout the world (with the exception of most European countries, the result of eradication programs), imposing significant economic costs on the livestock industry [5]. While not a natural host, it is also possible to experimentally infect sheep with BLV and in contrast to the situation in cattle, all infected sheep generally develop B cell leukemia/lymphoma, about 20 months post infection [6].

BLV infects B-cells where it integrates into the genome. Following an early and transient phase of horizontal replicative dissemination, BLV primarily proliferates via polyclonal expansion, producing many long lived clones that can be tracked via their proviral integration site [7,8]. In a subset of infected individuals, for unknown reasons, one of these clones eventually begins to expand, producing an aggressive neoplasm that accumulates in the blood (B-cell leukemia) and/or organs (B-cell lymphoma). In both BLV and HTLV-1 the Tax protein has been seen as the principal actor in oncogenesis, especially as it is capable of driving cellular transformation [9,10]. However, the lack of Tax expression in the majority of ATLs [11] and BLV induced tumors [12,13], points to a more diverse cast of actors beyond Tax. In the case of HTLV-1 it has been evident for the last decade that the antisense product HTLV-1 basic leucine zipper factor *(HBZ)* plays a central role in the process of oncogenesis as it is found in 100% of ATL cases examined [14,15]. A large number of roles have been attributed to HBZ, with its inhibition of Tax mediated viral transcription postulated as a vital part of the viruses immune evasion strategy [11,16,17]. Intriguingly, in addition to the protein encoded by *HBZ,* the *HBZ* mRNA has been implicated in supporting proliferation of ATL cells [18,19]. In the case of BLV, no equivalent antisense transcript has been described, although it has recently been reported by us and others that the provirus encodes a cluster of non-canonical RNA polymerase III transcribed microRNAs on the positive strand. These viral microRNAs represent ∽40% of the microRNA pool in transformed B cells [20,21], and like the case of HBZ in HTLV-1, they are expressed in all BLV induced tumors examined to date, pointing to a vital role in the life cycle of the virus and cellular transformation [20].

The close relationship between BLV and HTLV-1 in genome organization and pathogenesis makes BLV an attractive model for investigating deltaretrovirus induced cancer. To further our understanding of the transcriptional landscape of BLV infected cells and to build on our previous work looking at small noncoding RNAs we carried out high throughput RNA sequencing (RNA-seq) of total RNA from ovine and bovine BLV infected cells. Surprisingly, we observed a large number of reads mapping back to the BLV genome and found that these reads were the products of previously unidentified BLV antisense transcripts originating in the BLV 3’LTR. We present evidence that, like the case of HBZ, BLV antisense transcription appears to be a consistent feature of BLV infections. We also find that one of the transcripts is cleaved by the BLV microRNAs and discuss the possible role these transcripts play in the life cycle of BLV.

## Results

### Identification of two antisense transcripts encoded by BLV via deep sequencing

Previous work carried out by us and others has identified a cluster of ten microRNAs expressed from the BLV provirus genome utilizing a non-canonical microRNA biogenesis pathway involving RNA polymerase III [20,21]. In order to better understand the pattern of RNA transcription in BLV infections we expanded our sequencing efforts beyond small RNAs and sequenced stranded RNA libraries produced from a variety of BLV infected samples. In total 51 different samples were sequenced, including the ovine B cell lines YR2 & L267 derived from BLV induced primary tumors, PBMCs from asymptomatic pre-leukemic BLV infected sheep and BLV induced tumors isolated from both cattle and sheep. With the exception of the BLV microRNAs it has generally been assumed that the BLV provirus is transcriptionally silent in transformed B cells. Viral transcripts and proteins originating from the 5’LTR are generally undetectable as a result of either genetic or epigenetic modifications of the tumor provirus [13,22,23]. Stranded RNA-seq revealed a similar pattern. Sense reads observed were predominantly derived from overlapping host transcripts rather than 5’LTR initiated viral transcription (Rosewick, et al submitted) and there was an absence of split reads corresponding to the spliced transcripts of the structural and regulatory genes, including *tax* (Fig. 1A & Table S1). The situation for the antisense coverage was significantly different, with a substantial number of antisense reads observed. Antisense read coverage was concentrated at the LTRs (as the LTRs are identical reads originating from the 3’LTR will also map to the 5’LTR and vice versa) and a ∽300 bp region just downstream of the last BLV microRNA (miR-B5-3p). Additionally, the presence of split reads (between base pairs 7217 and 8353 in the BLV genome) pointed to a spliced transcript (denoted as Antisense 1, AS1). A second set of split reads were observed between base pairs 2829 and 8353, indicating the presence of a second transcript (denoted as Antisense 2, AS2). These putative splice sites do not overlap those found in previously described sense transcripts originating from the BLV 5’LTR (Fig. 1A). Additionally, splice donor/acceptor sites matched the negative strand, pointing to the 3’LTR as the origin of these transcripts.

In the BLV cell lines YR2 and L267 the number of viral antisense reads counted per million reads (CPM) was 7.4 and 9.7 respectively, while the CPM for sense reads was 0.5 and 0.2 respectively. This pattern was consistent across all the ovine and bovine tumors examined (Table S1). In the 28 ovine samples, viral antisense CPM varied from 0.4 to 124.5 (average 19.5, median 3.5), while the CPM of viral sense reads varied from 0 to 19.3 (average 1.5, median 0.6). The 12 bovine samples produced a similar pattern, with the antisense CPM ranging from 0.6 to 11 (average 5.4, median 4.9), the sense CPM ranged from 0 to 11.5 (average 2.3, median 1).

PCR primers designed to span the putative splice sites of AS1 showed amplification with cDNA derived from the YR2/L267 tumor cell lines, ovine and bovine tumors with no amplification in ovine cDNA (Fig. 1B; Fig. S1). PCR with a primer upstream of the BLV microRNAs and a primer in the first AS exon also showed amplification in YR2/L267 and the ovine bovine tumors (although two tumors showed fainter bands), demonstrating that the AS1 transcript could extend beyond the BLV microRNA region (AS1-L). In the case of AS2, the cell lines YR2/L267 and the tumors showed amplification, with the exception of the bovine tumor T1345, which carries a 4.4 kb deletion that removes the second exon of AS2 [20] (Fig. S1.)

**Figure 1.**
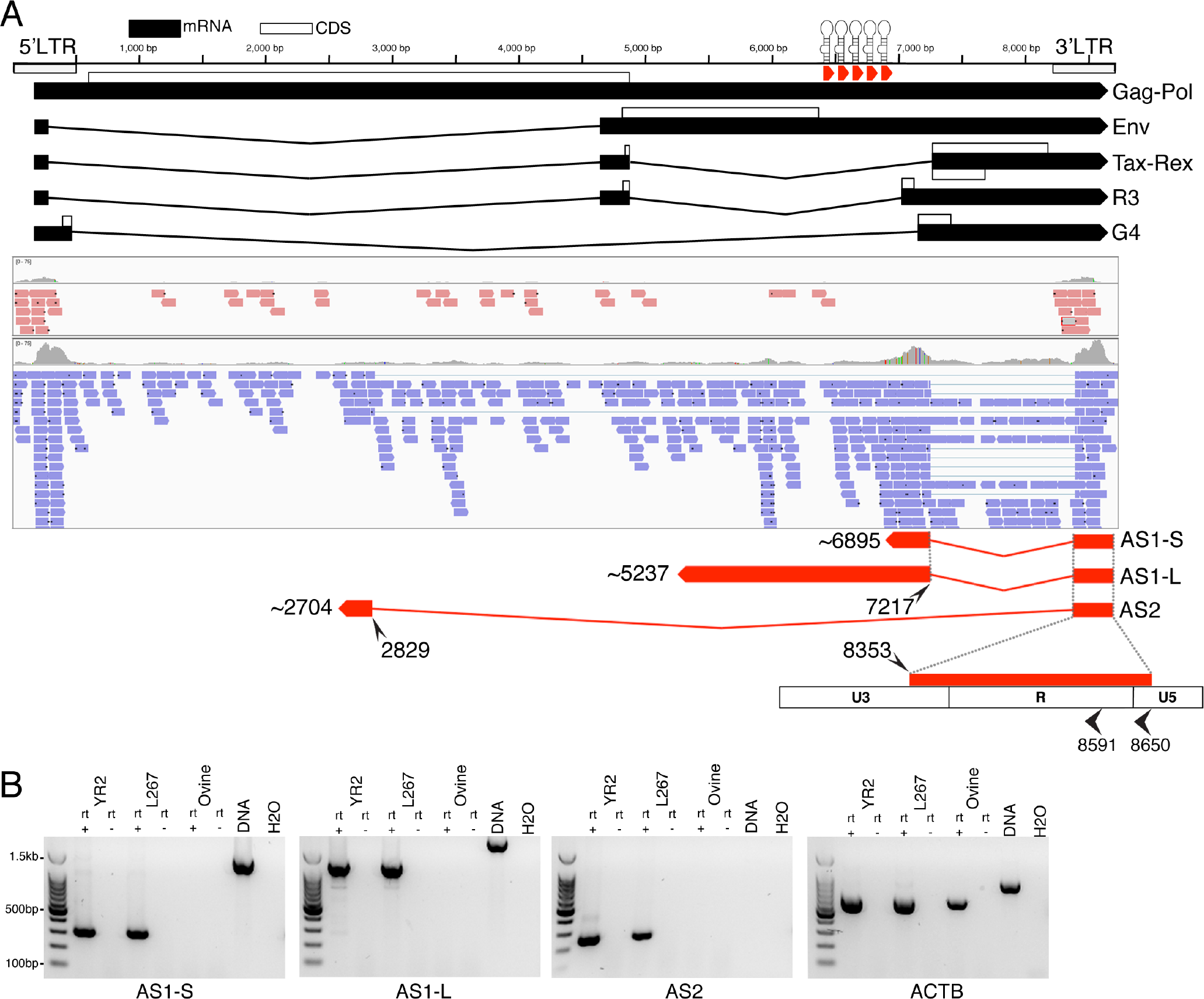
Identification of antisense transcription from the BLV provirus. **(A)** Ideogram showing location of the BLV sense transcripts originating in the 5’LTR as well as the BLV microRNAs. Below is shown a IGV [24] screen shot with the sense (red) and antisense (blue) reads from stranded RNA-seq of the L267 tumor B cell line. A limited number of sense read are observed and in the case of L267 these are derived primarily from the cellular transcript RASA3, rather than the viral 5’LTR (Rosewick et al submitted). In L267 and other samples examined antisense reads predominate. Based on observed antisense split reads and the results of 5’/3’ RACE three BLV antisense transcripts were identified. All three share the same exon 1, with the 5’end located primarily at either position 8591 or 8650. AS1-S ends as the transcript enters the BLV microRNA region and has a poly A tail. AS1-L extends beyond the BLV microRNA region and reads with a poly A tail can be observed at position ∽5237. Finally, AS2 has a small exon 2 (∽125bp) and no poly A tail was observed in the majority of samples examined. **(B)** Rt-PCR using the YR2 and L267 cell lines and PBMCs from an uninfected sheep. Control DNA was derived from YR2. Primers used span the splice sites for AS1-S\L and AS2. For AS1-L the reverse primer was placed upstream of the BLV microRNAs. Primers for Beta-actin (ACTB) were included as a positive control. (Rt+ reverse transcriptase positive, Rt-reverse transcriptase negative, base pair coordinates correspond to NCBI Accession: KT122858)

RNA sequencing of the YR2LTaxSN and L267LTaxSN cell lines [13], which constitutively express 5’LTR dependent viral transcripts and proteins, in addition to the RNA pol III dependent microRNAs, also showed the presence of antisense transcription (Fig. S2). The antisense CPM observed in these samples fell within the range seen for BLV samples lacking sense transcription (Table S1), suggesting that transcription from the 5’ and 3’ LTRs of BLV, as well as from the RNA pol III promoters, are not mutually exclusive. In addition to BLV induced cell lines, we also carried out RNA-seq of PBMCs isolated from six asymptomatic BLV infected sheep 17 months post inoculation. The proviral load in these animals ranged from 5.3% to 40.6% (average 14.4%, median 9.3%). High throughput sequencing (HTS) mapping of proviral integration sites showed that in five of these animals the largest clone (defined by percentage of reads observed at each insertion point) did not exceed 5% (Fig. 2A). In the animal with the highest proviral load (40.6%) a single clone dominated, accounting for ∽87% of the reads (Fig. 2A). Antisense reads were observed in all of the samples and at substantially higher numbers than sense reads. Additionally, the number of antisense reads mirrored the proviral load (Fig. 2B & Table S1). It is worth noting that we also observed the consistent production of BLV microRNAs in these early stage samples (Fig. S3) and that microRNA levels were proportional to the proviral load (Table S1). Altogether, it is apparent that BLV antisense transcription, like BLV microRNA production, is a feature of both the pre-leukemic and leukemic stages of BLV infection.

**Figure 2.**
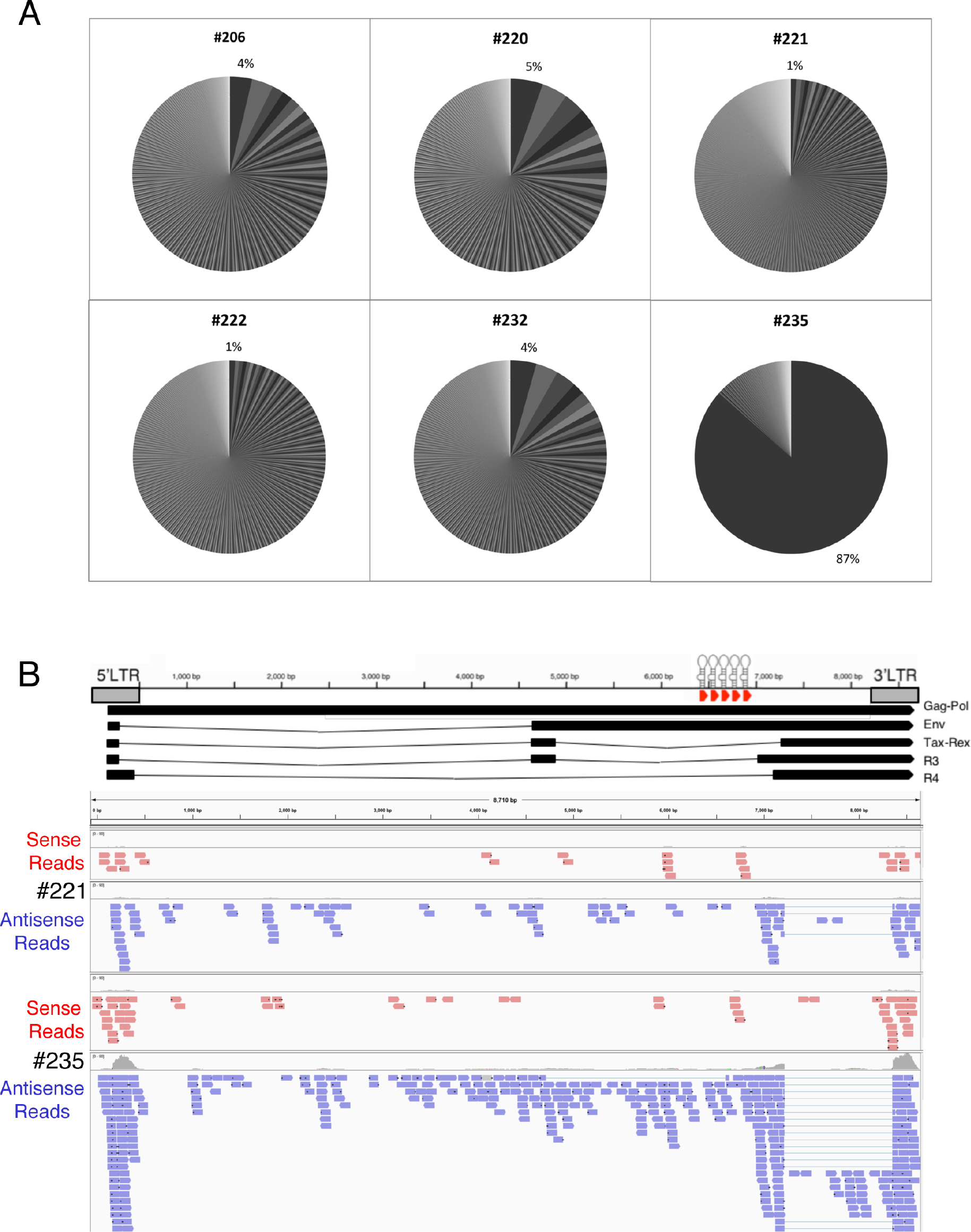
Antisense transcription is present at the polyclonal stage of infection. **(A)** Results of HTS mapping of provirus integration sites in six sheep 17 months post inoculation with the wild type pBLV344 provirus. Each slice of the pie graph represents the percentage of reads attributable to a single provirus insertion site. The percentage attributed to the largest clone is indicated. The first five animals did not possess any dominant clone, rather they show a large number of clones of low abundance (∽1000), with each accounting for a small percentage of the reads. The exception was animal &#x0023;235 where a single integration site accounted for 87% of the reads observed, signaling the expansion of the tumor clone. **(B)** Ideogram of BLV provirus with screen shots from IGV showing RNA-seq sense and antisense reads from the animals &#x0023;221 (polyclonal) and &#x0023;235 (tumor clone expanded).

### Identification of 5’/3’ ends

In order to determine the 5’ and 3’ ends of both antisense transcripts rapid amplification of cDNA ends (RACE) was carried out in combination with high throughput sequencing. For 5’RACE, total RNA from the cell line YR2 was used as template and mapping of the resultant reads showed that 42% of AS1 and 46% of AS2 transcripts started in U5 close to the R boundary at position 8650 (Fig. 1). A second site (8591) within the R region was also frequently utilized by both AS1 (27%) and AS2 (21%) (Fig. 1). Other less frequently used start sites are listed in (Table S2).

Our approach to 3’RACE was slightly different from traditional methods as we did not attempt to pick a transcript specific primer close to our assumed 3’ end. Rather we used a primer in the first exon of AS1/2 to produce nearly full length cDNAs, which were then processed into libraries for high throughput sequencing. This approach was taken to increase the chance of identifying alternative poly A tails in the transcripts. Libraries were produced using total RNA from the cell lines YR2, L267 and FLK, three ovine and four bovine tumors, as well as six BLV infected sheep 17 months post inoculation. When mapped back to the BLV genome, the products of 3’ RACE were mainly clustered in a region overlapping with BLV miR-B5-3p (Fig. S4). A canonical AAUAAA polyadenylation signal sequence (PAS) was found at position 6913-6918 immediately adjacent to the end of miR-B5-3p, however no GU rich consensus sequence was obvious (Fig. S5A). The observed poly A sequences were not confined to a single position but clustered at various points in a ∽60bp region between 6842-6910. This region also corresponds with a position where the antisense read coverage drops, indicating that the majority of BLV antisense transcripts utilize this poly A site, resulting in a short transcript (AS1-S) of approximately ∽600bp (Fig. 1 & Fig. S4). In all the samples, a number of reads were observed beyond this point, with a second cluster of poly A sequences observed around position 6171. No PAS was observed adjacent to this position and the presence of an adenine rich region overlapping with the poly A sequences suggests that this region may have served as the priming site for our oligo dT primer. As a consequence, it does not appear to be a legitimate poly A tail. A second canonical AAUAAA (PAS) was found at position 5257-5262 (again without an obvious GU rich consensus sequence). When 3’RACE was carried out with a primer adjacent to this PAS and YR2 as template, reads were observed 20bp upstream (Fig. S5B). Reads showing evidence for this poly A tail were also observed in five of the libraries produced using a primer extending from the first AS exon. An additional 3’RACE library, using YR2 RNA as template, was sequenced on an Oxford Nanopore MinION. The long reads from this instrument showed a number of transcripts extending up to the region around position ∽5237 (Fig. S6A). The presence of a longer version of AS1 is also supported by PCR with the forward primer in the first AS1 exon and a reverse primer upstream of the BLV microRNAs, which produces robust amplification in the majority of BLV tumors examined (Fig. 1B & Fig. S1). Together these results indicate that a portion of the AS1 transcripts do extend into and beyond the BLV microRNA region, creating a long AS1 transcript (AS1-L). While some AS1-L transcripts appear to be polyadenylated at position ∽5237 the frequency appears low and in the MinION reads many of the AS1-L reads reveal fusions with the host genome (Fig. S6B). Additionally, it should be noted that the bovine tumor T1345 contains a large internal deletion that includes the PAS at position 5257-5262 (includes the last ∽100bp of AS1-L, assuming poly A tail at position ∽5237) suggesting that at least in tumors, this portion of the transcript is dispensable.

In the case of AS2, the majority of the samples showed no evidence of a poly A tail, rather the AS2 transcript ended abruptly at position 2704. Closer examination of the sequences showed the transcript to have undergone splicing with the host genome, creating a BLV-host fusion transcript. In each tumor examined, the sequence forming the fusion with AS2 corresponded to the region of the host genome where the provirus was inserted (Fig. S5C). In the case of L267, in addition to BLV-host fusion transcripts, a small number of AS2 reads were observed with appetent poly A tails (Fig. S5C). No PAS was observed in the vicinity and the position of the poly A varied over a ∽220 bp region with the majority slightly upstream of the splice site utilized in the formation of the BLV-host fusion transcript.

### BLV 3’LTR promoter activity and regulatory sequences affecting antisense transcription

The capacity of BLV to drive sense transcription from the 5’LTR has been investigated in some detail over the years, however the promoter activity and landscape of regulatory motifs capable of driving antisense expression from the 3’LTR remained unexplored. To confirm that the 3’LTR was capable of driving expression of an antisense transcript, we inserted the BLV LTR upstream of a luciferase construct in both the 5’ and 3’ orientations (Fig. 3A). In the 3’ orientation, luciferase activity was significantly higher than that seen in the 5’ orientation (Fig. 3B), consistent with BLV 3’LTR antisense promoter activity. The majority of regulatory motifs identified in the BLV 5’ LTR are concentrated in the U3 region. Removal of the first 167bp of U3 in the 3’LTR construct (3’Δ) (Fig. 3A), rather than inhibiting transcription, resulted in significantly increased luciferase activity (Fig. 3B). As expected, expression of Tax, the potent viral transactivator, resulted in a dramatic increase of luciferase activity with the LTR in the 5’ orientation (Fig. 3C), while in the 3’ orientation Tax caused a significant reduction in luciferase activity (Fig. 3D).

It has previously been reported that in addition to the many regulatory regions in the U3 region there are also an Interferon Regulatory Factor (IRF) binding site in U5 [25] and a E-Box in the R region (labeled as DAS: downstream activator sequence) [26]. In order to test if these motifs played a role in the 3’LTR antisense promoter activity, we carried out site directed mutagenesis on both sites in the BLV LTR constructs. Disruption of the DAS motif caused a significant drop in luciferase activity for both the 5’LTR and 3’LTR constructs. In the case of the IRF motif, a modest although non significant reduction was seen in both constructs (Fig. 3E). Finally, disruption of the IRF and DAS motifs in the construct carrying the 167bp deletion in U3 caused a significant reduction in luciferase activity for both motifs (Fig. 3E). Taken together these results show the 3’LTR is capable of driving antisense transcription in the BLV provirus and that the IRF and DAS motifs appear to play a role in both sense and antisense transcription.

**Figure 3.**
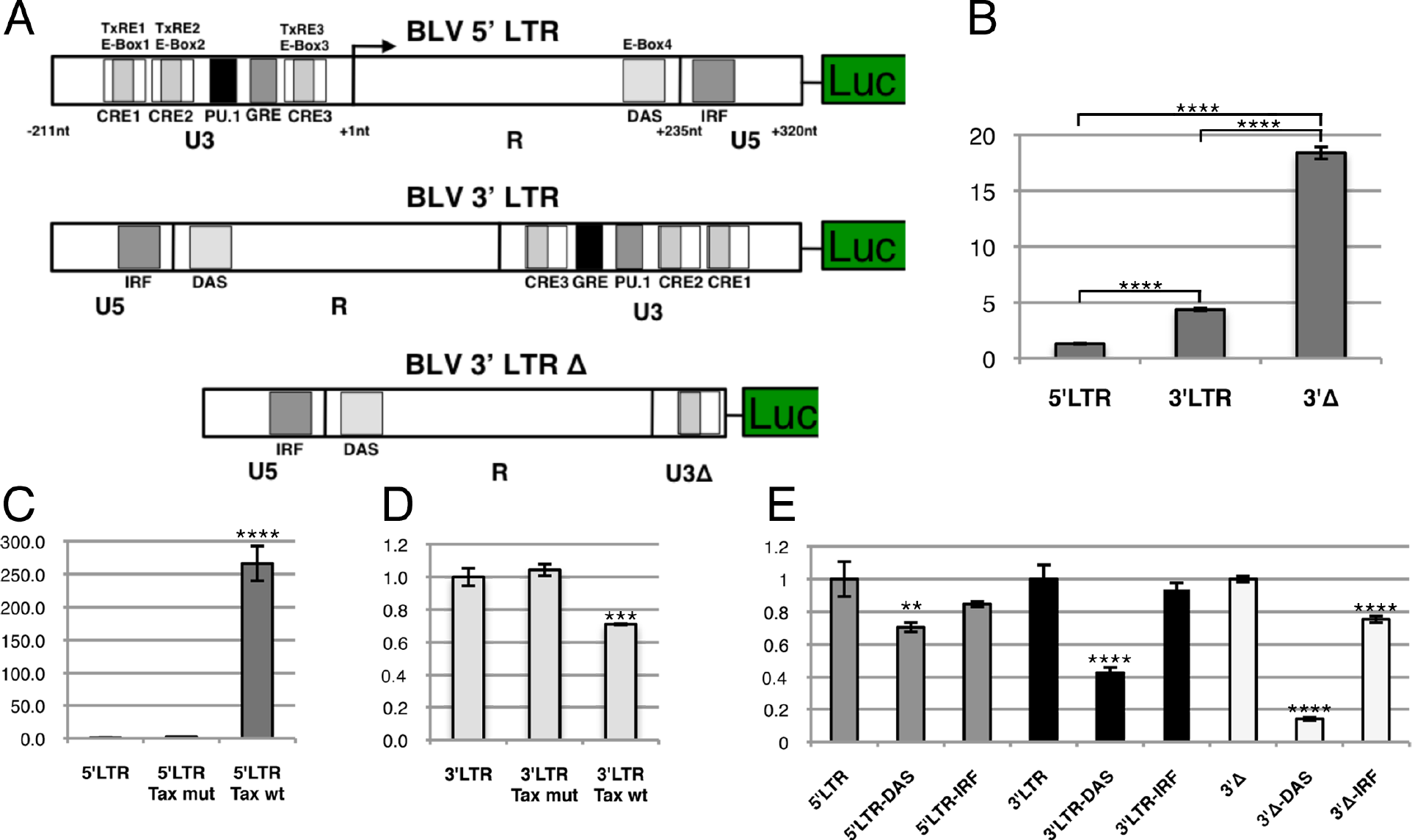
BLV 3’LTR antisense promoter activity and regulatory sequences affecting transcription. **(A)** Ideogram showing the three constructs used in luciferase assays, the first with the orientation of the BLV LTR matching that in the 5’LTR, the second matching the 3’LTR and the third in the 3’LTR orientation but with the majority of the U3 region deleted (3’Δ). Transcription factor binding sites from [27]. **(B)** Luciferase levels for the 5’LTR, 3’LTR and 3’Δ constructs. Comparison of the three constructs showed a significant difference between each. **(C)** Tax is a powerful transactivator of the 5’LTR. Transfection was carried out with the 5’LTR construct alone, with the 5’LTR construct in conjunction with a Tax expressing plasmid (5’LTR Tax wt) and finally with the 5’LTR construct in conjunction with a mutated version of Tax incapable of driving transactivation (5’LTR Tax mut). The addition of Tax caused a significant increase in luciferase levels. **(D)** Comparison of luciferase levels in the 3’LTR construct following the addition of Tax. Transfection with the 3’LTR construct (3’LTR) and co-transfection with Tax (3’LTR Tax wt) or with the mutated Tax (3’LTR Tax mut). The addition of wt Tax caused a significant drop in luciferase levels. **(E)** The IRF and DAS (E-Box4) transcription factor binding sites present in U5 and R respectively, were mutated in each of the constructs. In the 5’LTR construct, mutation of IRF caused a modest but non significant drop in expression. Mutating DAS causes a more pronounced and statistically significant drop, an observation in line with previous reports. In the case of the 3’LTR, the construct with a mutation in IRF showed a modest, but non significant drop in expression, while mutating DAS caused a more dramatic and significant drop, reducing expression levels by more than 50%. Finally, in the 3’Δ construct, mutating IRF & DAS caused significant drops in luciferase levels, with the mutation in DAS reduced expression by over 80%. (Luciferase levels are scaled to the left most construct in B,C and D. In E they were scaled to the appropriate wild type construct. Statistical significance was determined via Tukey's multiple comparisons test, **p-value < 0.01, ***p-value < 0.001 ****p-value < 0.0001)

### Coding potential

Having confirmed the promoter activity of the 3’LTR and estimated the approximate scope of the BLV antisense transcripts, we next examined them for coding potential using the Coding Potential Assessment Tool (CPAT) [28]. The limited number of annotated bovine long non-coding RNAs (lncRNAs) prevented the development of a robust bovine specific model to calculate coding probability. As a result, human and mouse CPAT models were utilized. In addition to the BLV antisense transcripts, a number of known BLV and HTLV-1 protein coding genes and 220 bovine lncRNAs were analyzed. Using CPAT the known protein coding genes, including the HTLV-1 antisense gene *HBZ,* were all assigned a coding probability greater than 0.98 using the human model, and greater than 0.91 in the mouse model. In contrast, >90% of the bovine lncRNAs were assigned a coding probability less than 0.25 in both models (Table S3). AS2 coding probability was low (0.003 human model, 0.022 mouse model), allowing us to classify it as a lncRNA. However, AS1-S/L were assigned an intermediate, ambiguous coding probability of ∽0.46 in the human and ∽0.39 in the mouse model, with a potential open reading frame (ORF) of 264bp (shared by both AS1-S & AS1-L). The small putative peptide sequence did not show any significant matches when used to search the non-redundant protein sequence (nr) database with BLASTP and the protein families database [29].

We next examined patterns of nucleotide variation in the region containing the putative peptide. Using our RNA-seq data, we extracted the sequence of the putative AS1-S/L ORF from each of our bovine and ovine tumors and supplemented these with six publicly available whole genome BLV sequences from NCBI. AS1-S/L was characterized by a nucleotide diversity (average difference per nucleotide site for all pairwise comparisons of available sequences) of 0.0149. For comparison Tax, known to be essential, had a nucleotide diversity of 0.0106. Throughout the putative ORF, the majority of the nucleotide diversity of AS1-S/L, like that of Tax, was concentrated in the less constrained 3^rd^ codon position of the negative strand (0.034 AS1-S/L & Tax 0.025 respectively) (Fig. S7 & Table S4). It should be noted though that for just over 2/3 of the sequence, the potential AS1-S/L ORF overlaps with parts of the R3 and G4 ORFs on the sense strand. Additionally, the 3^rd^ codon position of G4 on the sense corresponds to the 3^rd^ codon position of the potential AS1-S/L ORF on the negative strand (Fig. S7). Nevertheless, this pattern of nucleotide diversity was not confined to the overlapping regions and extends into the region containing no overlapping BLV sense ORF (no overlap 0.0236 vs no overlap 3^rd^ 0.0521; Fig. S7 & Table S4). As a consequence the evidence for coding potential in AS1-S/L is suggestive, but not definitive, making it best to classify AS1-S/L as transcripts of unknown coding potential (TUCP) at the present moment [30].

### Sub-cellular localization of BLV antisense transcripts

Given the ambiguous coding potential of the BLV antisense transcripts we next sought to determine their location in the cell to help us understand potential functions. This was achieved by carrying out high throughput sequencing of total and small RNA libraries using cytoplasmic and nuclear enriched RNA fractions from the YR2 cell line. The resultant small RNA libraries showed the BLV microRNAs (like the cellular microRNAs) to be enriched in the cytoplasmic fraction (Fig. 4). In the case of the total RNA libraries, the BLV antisense reads (all antisense reads were considered together) were enriched in the nuclear fraction (Fig. 4). Absolute quantification via real-time PCR using primers spanning the splice site of AS1-S/L and AS2 also showed both transcripts to be enriched in the nuclear RNA fraction (Fig. 4).

**Figure 4.**
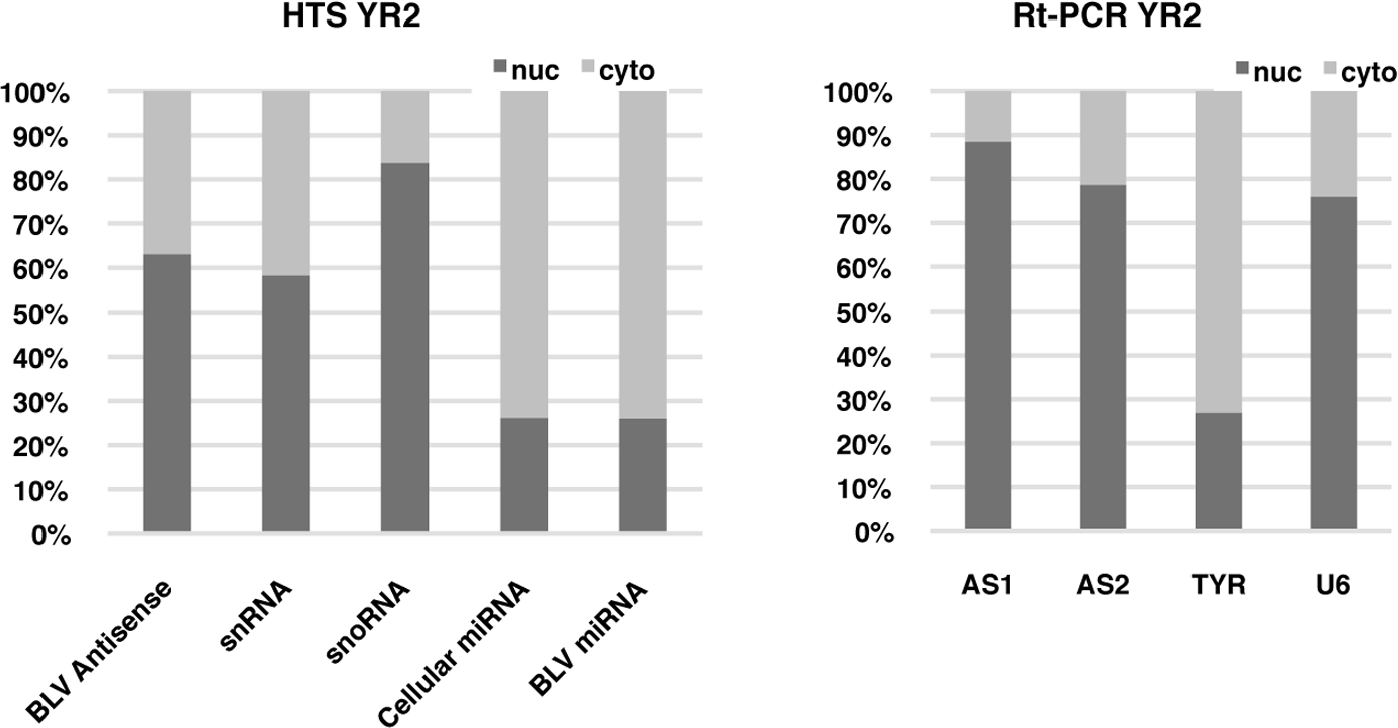
The YR2 cell line was fractionated to produce cytoplasmic and nuclear enriched pools of RNA that were utilized in the production of total RNA and small RNA libraries for high throughput sequencing (HTS). Read count for each library was normalized and enrichment in the cytoplasm/nucleus calculated for the different classes of transcript in YR2. The HTS results were supplemented with RT-PCR using primers for the antisense BLV transcripts AS1-S/L and AS2. Primers for TYR (cytoplasmic) and U6 (nuclear) were used to confirm fractionation efficacy. Both approaches show that the BLV antisense transcript has a primarily nuclear localization. (snoRNA = small nucleolar RNA; snRNA = small nuclear RNA)

### BLV microRNA guided cleavage of AS1-L

The majority of the AS1 transcripts is polyadenylated at a position just upstream of the BLV microRNAs (AS1-S). However, we also observed that approximately ∽19% of the antisense transcripts extended beyond this point (Fig. S4, S5 & S6). These AS1-L transcripts overlap the region encoding the BLV microRNAs and as the transcripts are derived from opposite strands there will be perfect complementarity between them. Consequently, the AS1-L transcript should be sliced via the action of RISC (RNA-induced silencing complex), with cleavage occurring between the 10^th^ and 11^th^ base pair position of the mature microRNA [31]. To identify the products of cleavage, a modified version of RNA-ligase-mediated (RLM) 5’ RACE [32] was employed and combined with high throughput sequencing to determine the precise position of the 5’ end of each read in the BLV genome. Using RNA from the YR2 cell line, we found that ∽66% of the reads mapping to the ∽550bp BLV microRNA region had a 5’ end that fell between the 10^th^ and 11^th^ base pair of one of the BLV microRNAs. The majority of the cleavage appears to be driven by five of the BLV microRNAs. Of these, miR-B1-3p was responsible for 35.7% of the reads, miR-B2-5p 3.4%, miR-B2-3p 7.5%, miR-B4-3p 16.2% and miR-B5-3p 2.9% (Fig. 5 & Table 1). We also produced a library using total RNA extracted from the PBMCs of a sheep 19 months post infection and found that ∽26% of the reads mapping to the BLV microRNA region showed evidence of cleavage mediated by the BLV microRNAs (Table 1).

To examine if this cleavage was occurring in the cytoplasm or nucleus, we produced nuclear and cytoplasmic enriched RNA pools from YR2 and carried out the same modified 5’RACE and sequencing. There were more than double the number of reads showing evidence of cleavage mediated by the BLV microRNAs in the cytoplasmic fraction (∽54%) when compared to the nuclear fraction (17%), indicating RISC mediated cleavage in the cytoplasm (Fig. 5 & Table 1). The percentage of reads mapping to the BLV microRNA region was similar within each fraction (∽56% nuclear, ∽57% cytoplasmic).

**Figure 5.**
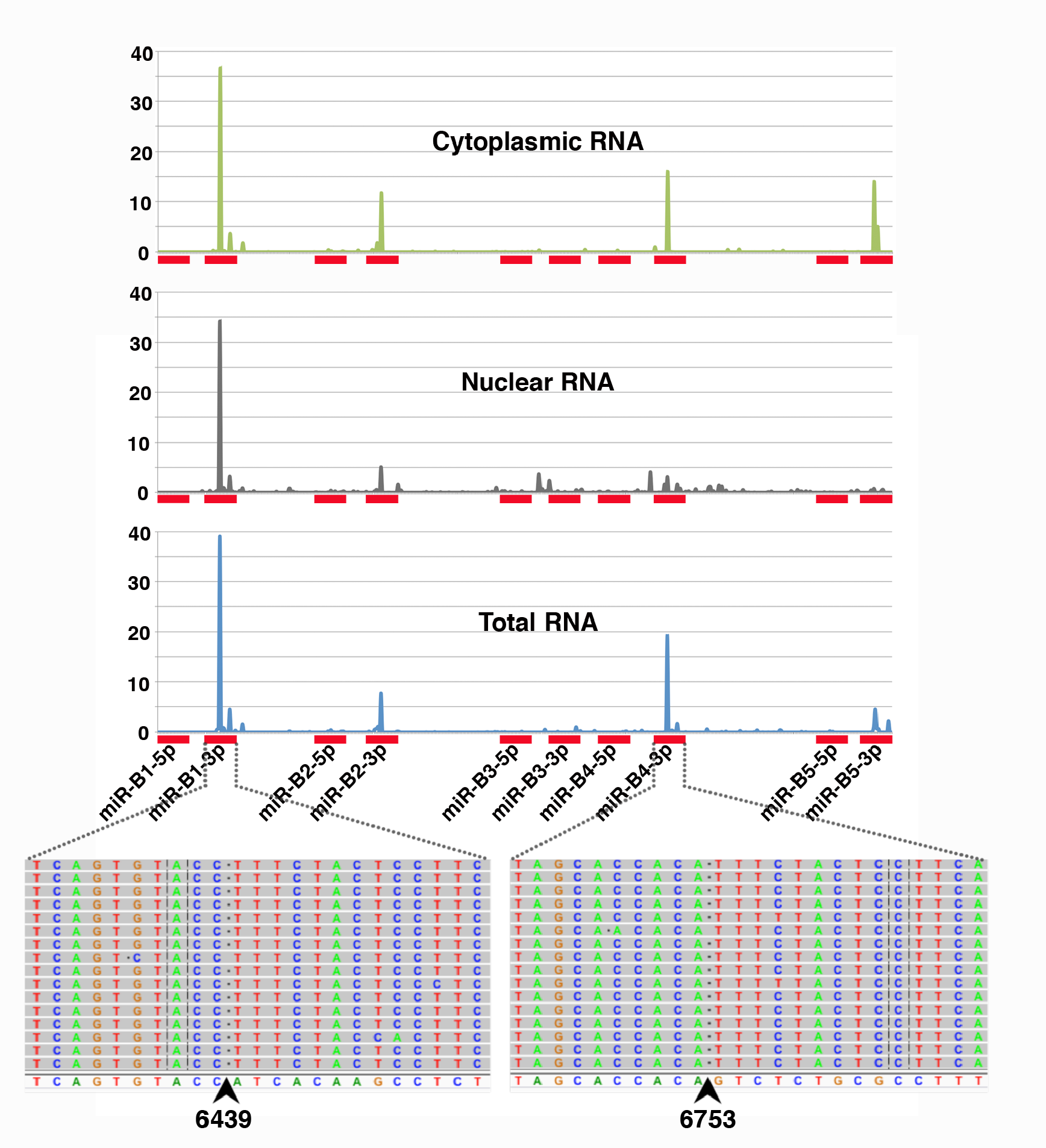
BLV microRNA mediated cleavage of the AS1-L transcript. Shown on the X axis is the region of the BLV genome containing the BLV microRNAs (denoted by red rectangles). Graphed above is the percentage of high throughput sequencing reads showing evidence of cleavage at each base position. Libraries were prepared from total YR2 RNA and YR2 cells that were fractionated into cytoplasmic and nuclear enriched fractions. Below the X axis are screen shots from IGV showing the precise base pair position of cleavage for the peaks observed at miR-B1-3p and miR-B4-3p. The reference sequence is shown at the bottom and the individual reads above. The first 10 bp of the reads match the reference sequence, while the sequence to the right of the cleavage point (arrow) corresponds to the RNA oligo ligated to the free 5’ end. Sites of cleavage were enriched between bp 10 & 11 of a subset of the mature microRNAs. The frequency of reads showing evidence of cleavage was increased in the cytoplasmic fraction.

**Table 1.** Percentage of reads in the BLV microRNA region that show evidence of cleavage mediated by each microRNA (between the 10 and 11bp of the mature microRNA)

|  | YR2 Total | YR2 nuc | YR2 cyto | #235 |
| --- | --- | --- | --- | --- |
| miR1-5p | 0 | 0 | 0 | 0 |
| miR1-3p | 35.71 | 11.91 | 26.21 | 19.68 |
| miR2-5p | 3.38 | 2.00 | 3.97 | 2.33 |
| miR2-3p | 7.55 | 1.59 | 8.50 | 0.56 |
| miR3-5p | 0.22 | 0.00 | 0.00 | 0.00 |
| miR3-3p | 0.00 | 0.00 | 0.00 | 0.00 |
| miR4-5p | 0.00 | 0.00 | 0.00 | 0.00 |
| miR4-3p | 16.16 | 1.22 | 4.68 | 3.09 |
| miR5-5p | 0.23 | 0.03 | 0.02 | 0.00 |
| miR5-3p | 2.90 | 0.34 | 10.61 | 0.01 |
| Total % | 66.14 | 17.08 | 53.99 | 25.67 |

### Deleting and inverting the BLV microRNAs

Given the observed cleavage of AS1-L in the cytoplasm we next sought to examine if the absence of BLV microRNAs would have any effect on the relative fraction of AS1-S/L produced. To achieve this, we designed two altered BLV proviruses. In the first, the BLV microRNAs were removed, while in the second they were inverted to prevent cleavage of AS1-L (in both cases the PAS just upstream of BLV miR-B5-3p was also removed, Fig. S8A). These proviruses were transfected into HeLa cells, followed by total RNA extraction and 3’RACE with the forward primer in the first common AS exon followed by high throughput sequencing. Like the situation in tumors, the WT provirus 3’RACE library showed a rapid coverage drop off in the region containing the BLV microRNAs (Fig. S8A). In contrast, for the deleted provirus the level of coverage remained relatively consistent across the length of AS1-L, while in the inverted provirus, there was a gradual decrease in coverage along the length of AS1-L (Fig. S8A). As a result, removal or inversion of the BLV microRNAs and the adjacent PAS appears to increase the fraction of AS1-L transcripts present.

The increased fraction of AS1-L transcripts could be explained in two ways. Firstly, the RNA pol III sitting over the microRNAs may inhibit RNA pol II progression while the PAS adjacent to BLV miR-B5-3p microRNA mediates polyadenylation, thereby forming AS1-S. By removing these elements, AS1-L levels increase. Secondly, by removing the microRNAs we prevent the cleavage of AS1-L in the cytoplasm, increasing the apparent number of AS1-L reads and causing their accumulation in the cytoplasm. In an attempt to distinguish between these different scenarios, we fractionated transfected cells in into nuclear and cytoplasm enriched RNA pools. Real-time PCR with primers spanning the splice sites for AS1-S/L, AS2 and BLV sense transcripts did not show differences in the sub-cellular distribution of transcripts for the three proviruses (Fig. S8B). As a result, it does not appear that the increased fraction of AS1-L transcripts observed is a consequence of eliminating BLV microRNA mediated slicing of AS1-L, rather it suggests the first scenario involving RNA pol II and pol III collisions.

### Knock down of AS1 with locked nucleic acids (LNAs)

In a final effort to gain an insight into the role of the BLV antisense transcripts we sought to knock down their expression in YR2 and L267. Both cell lines, derived from ovine primary B lymphoid tumors, were previously shown to carry a transcriptionally silent provirus impaired in 5’LTR dependent transcription of protein coding genes [23]. As these B cell lines are refractory to transfection, we utilized locked nucleic acid antisense oligos, introduced via unassisted uptake [33]. A mix of three LNAs, targeting different parts of AS1 were used to treat both cell lines at concentrations of 5uM and 10uM. In addition, the cell lines were treated with 10μM of control LNAs and with a mock treatment of H20. Analysis of the real-time PCR results with the Kruskal-Wallis test (One-way ANOVA on ranks) showed a significant difference between the four treatments for AS1-S/L (p value = 0.0192). Dunn's multiple comparisons test was then used to examine differences between the individual samples. For the 5μM concentration, the observed drop in expression (20%) failed to reach statistical significance in any comparison. In the case of the 10μM concentration, the more pronounced drop (>50%) between the control and AS specific LNAs was significant (p value = 0.0347). In the case of AS2 a significant difference was observed between the treatments (Kruskal-Wallis test, p value = 0.0208), however Dunn's multiple comparisons test failed to show any difference between individual samples (Fig. 6). The real time assay for Tax failed to show amplification or was indistinguishable from the background in all the YR2 and L267 treatments (Tax expressing cells showed robust amplification with an average Ct of 21.5). Indicating that knock down of AS1 does not cause a detectable reactivation of transcription from the 5’LTR. Finally, there was no apparent difference in the viable cell counts in the treated and control cells at the end of the experiment (Fig S9).

**Figure 6.**
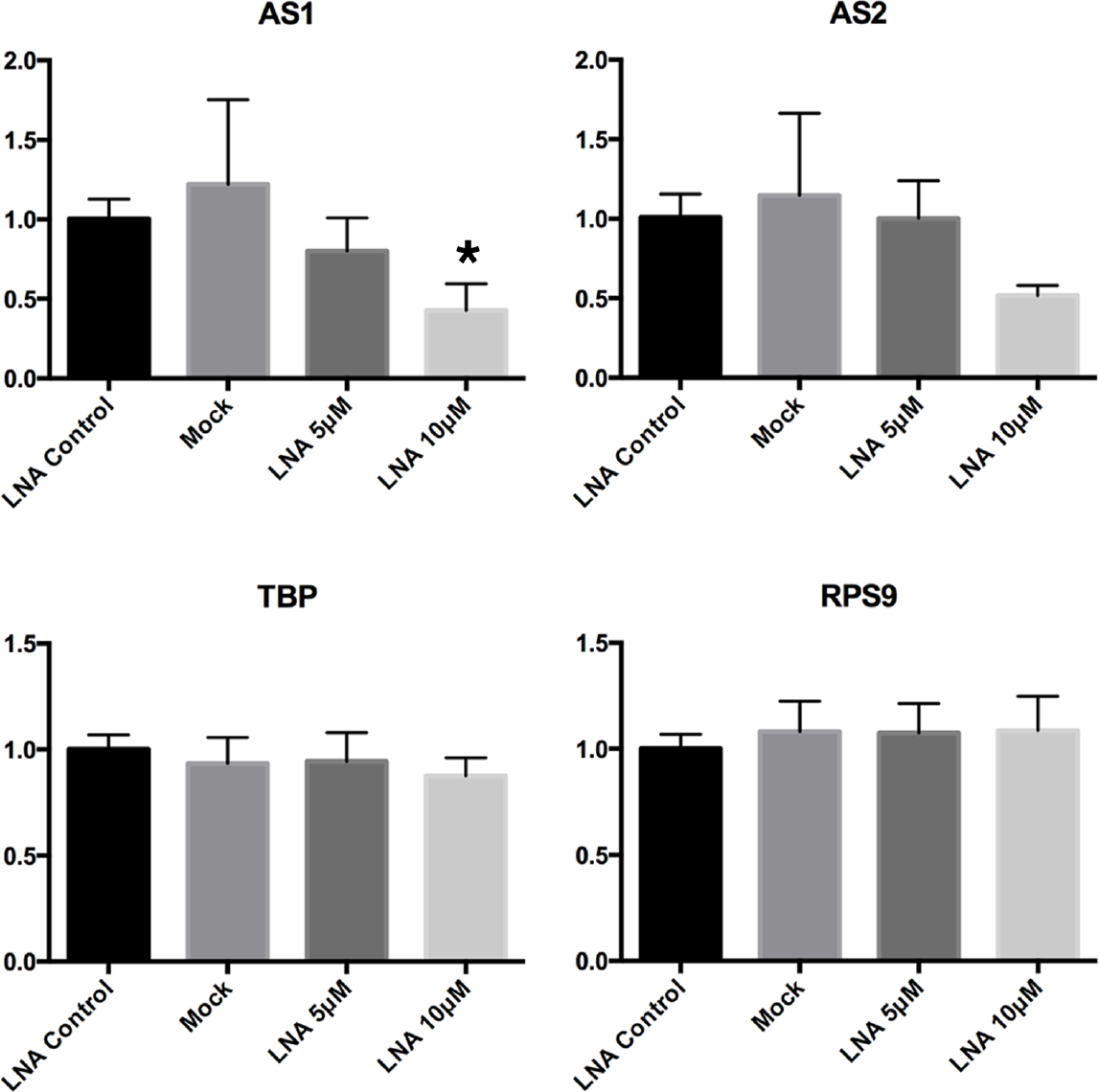
Knock down of AS1 with locked nucleic acids. Three locked nucleic acids (LNAs) designed to target the common first exon of AS1/2, the AS1 splice junction and the second exon of AS1 were mixed and introduced via unassisted uptake at 5 |lM and 10 |lM concentrations, in duplicate to the YR2 and L267 BLV cell lines. Scrambled LNAs at a concentration of 10 |lM and a mock treatment were also carried out. Following 5 days of incubation, RNA was extracted and real time PCR carried out with assays for AS1, AS2, Tax and the house keeping genes RSP9 and TBP. Fold changes observed were scaled to the LNA control samples, which were set to 1. The Kruskal-Wallis test showed a significant difference between the treatments in AS1 (p-value = 0.0192), with Dunn's multiple comparisons test showing a significant difference in LNA Control vs. LNA 10|lM (adjusted *p-value < 0.05). For AS2 Kruskal-Wallis test showed a significant difference between the groups, but Dunn's multiple comparisons test did not show any significance between the individual treatments. Tax expression was not detected in the treated or control YR2 and L267 cells and was not graphed. (Values are means ± SD combined for YR2 and L267.)

## Discussion

The identification of BLV antisense transcription is in hindsight somewhat unsurprising. It has been known for over a decade that BLV's close relative HTLV-1 encodes an antisense transcript [17] and like HTLV-1, the BLV 3’LTR remains unmethylated in tumors [23], begging the question of a BLV antisense transcript. Prior to the adoption of RNA-seq, identifying these transcripts required a certain amount of luck in the placement of primers/probes for Rt-PCR/Northern blotting. However, as can be seen from Fig. 1 the use of RNA-seq, especially in combination with stranded libraries, made the presence of antisense reads in BLV obvious. The consistent pattern of antisense transcription across all the BLV samples sequenced to date strongly suggests that these transcripts fulfill an essential function in the biology of BLV and leukemogenesis. Furthermore, in the vast majority of samples little or no evidence of sense transcription was observed, again highlighting that Tax is not required for the maintenance of cellular transformation, mirroring results recently obtained in HTLV-1 [15].

Our initial observations using BLV induced tumors showed consistent antisense transcription, but little or no sense transcription. This lead us to question if antisense/sense transcription were mutually exclusive. We addressed this question in a number of ways. Firstly, we carried out RNA-seq of the cell lines YR2_LTaxSN_ and L267_LTaxSN_, which both constitutively express transcripts from the 5’LTR and viral proteins. Antisense CPM for both (3.6 and 29.4 respectively) fall within the range observed for ovine and bovine tumors while showing sense CPM orders of magnitude higher than that observed in tumors (1,001 and 81,482 respectively) (Table S1). The pattern of antisense splicing also remains consistent in these cell lines (Fig. S1). Furthermore sequencing of a 3’RACE library from the FLK cell line (a fetal lamb kidney cell line infected with BLV that constitutively expresses BLV sense transcripts/proteins) showed a distribution of reads comparable to that observed in the ovine and bovine tumors (Fig. S4). Based on these observations, it seems that BLV 5’LTR dependent sense expression, in addition to being independent of BLV microRNA expression, does not preclude antisense transcription or create obvious changes in the pattern of AS splicing. However, as these observations are based on in vitro samples, the situation in vivo may be different.

Having found consistent antisense transcription in tumors, we next sought to determine if the same was true in pre-leukemic clones. Stranded RNA-seq of PBMCs from six asymptomatic BLV infected sheep revealed the presence of BLV antisense transcription and showed that antisense exceeded sense transcription in all cases (Table S1). Additionally, sequencing of small RNA libraries from three of these sheep showed evidence of robust BLV microRNA expression (Table S1). We could also be confident that the antisense/microRNA transcription in asymptomatic stages of infection was not driven by a single dominant clone, as the majority of the samples examined revealed the presence of multiple clones, with each clone representing only a small fraction of the overall proviral load (Fig. 2). Thus, BLV antisense transcripts, like BLV microRNAs, are present before the emergence of the tumor clone/clones and are produced by the majority of clones as antisense CPM increases in step with proviral load (Table S1).

The presence of BLV antisense transcription implies that the BLV LTR (like that of HTLV-1) is capable of driving transcription in both directions. Extensive work over the years has revealed a rich set of regulatory motifs bound by cellular transcription factors and Tax [26,27], regulating the BLV 5’LTR. As expected given the evidence of antisense transcription, we found that the BLV LTR was able to drive transcription in the orientation of the 3’LTR and with higher activity compared to the weak basal activity typical of the 5’LTR. The removal of the majority of the regulatory motifs in the U3 region increased the 3’LTR construct's activity, indicating that these motifs inhibit transcription originating in U5. While the majority of regulatory motifs identified in the BLV LTR are concentrated in the U3 region, a small number have also been identified in the R and U5 regions. Mutating the IRF motif in U5, while not significant in the 3’LTR construct did prove significant in the 3’Δ construct, suggesting a role for this motif in both the 5’ and 3’LTRs. In the case of the DAS (E-box4) found in the R region, when mutated, promoter activity was reduced dramatically in 3’LTR and 3’Δ constructs pointing to an important role in driving expression from the 3’LTR. However, as we are dealing with an artificial construct removed from the rest of the provirus, how closely these observation match the situation in the provirus remains to be seen. Nevertheless, it will be interesting to see if future work can tease out how the interplay between the transcription factors binding to the U3 and U5 regions of the LTR regulate transcription and how they shepherd transcription in the appropriate direction.

The Tax protein is a potent transactivator of the 5’LTR and as expected, the addition of Tax caused a dramatic increase in transcription for our 5’LTR luciferase construct. A similar pattern was not observed for the 3’LTR antisense promoter construct, rather transcription levels were reduced, indicating that Tax is not required for, and actually may have an antagonistic effect on BLV antisense transcription. Consistent with these results, we observed the constitutive production of AS transcripts in primary tumors, despite the complete absence or genetic inactivation of *Tax.* Tax is not required for maintaining malignancy and it is widely accepted that the absence of this immuno-dominant viral product in transformed cells plays a vital role in the virus immune evasion strategy. In contrast, the permanent production of both antisense transcripts and microRNAs, besides pointing to an essential function in tumorigenesis, indicate that these transcripts are incapable of eliciting an effective host immune response.

Having established the presence of the BLV antisense transcripts and determined that the 3’LTR was capable of driving its expression we sought to determine the approximate size of the transcripts. The use of alternate polyadenylation sites and the formation of fusion transcripts by the transcripts made precise demarcation of the 3’ends somewhat difficult. By using a combination of priming RACE in the first common exon of the antisense transcripts, high throughput sequencing and long reads we were able to show that the majority of AS1 is only ∽600bp in length, a much shorter transcript than *HBZ* (>2kb) in HTLV-1. This combined approach also showed that AS2 exon 2 was small (∽125bp) and generally forms fusion transcripts with the host genome. We also observed a longer version of the AS1 transcript (>2kb) that extended beyond the BLV microRNAs, however determining the end of this transcript was somewhat problematic due to biases in our 3’RACE PCR towards shorter fragments. To avoid this bias, 3’RACE with a primer adjacent to the canonical AAUAAA (PAS) at 5257-5262 was carried out in addition to the examination of long transcripts with the MinION. Both approaches showed that AS1-L reads could extend to this location. However the balance between the use of this poly A tail and the formation of fusions with the host genome remains unknown at the moment and will likely depend on the location of the provirus in the genome. The use of long read technologies such as the MinION will be particularly useful in this regard as full length sequences can unambiguously determine the structure of fusion transcripts.

The BLV AS1 transcript is found in a similar region of the proviral genome as that occupied by *HBZ* in HTLV-1, with the AS1-L variant producing a similar sized transcript. RNA-seq indicated that the majority of the AS1 reads take the form of the AS1-S variant, which terminates just before the BLV microRNA region, and is considerably smaller than *HBZ.* In the case of the AS1-L, there is an overlap with the BLV microRNAs leading to the cleavage of AS1-L via action of the RISC complex [34]. Our modified 5’RACE experiments confirmed this and showed that RISC mediated cleavage predominated in the cytoplasm (the fraction also enriched for the BLV microRNAs in YR2). The limited number of microRNAs involved in cleavage is intriguing as previous work has shown that all the tested BLV microRNAs associate with the RISC complex [20]. This may point to secondary structures in BLV AS1-L that inhibit cutting at certain positions, however the significance of this observation remains to be determined and more samples are required to see if this pattern is consistent. A further question that will require future work is the fraction of reads that are subjected to cleavage. The majority of the AS1 transcripts (AS1-S) do not contain the region complementary to the BLV microRNAs. Additionally, it appears that a majority of the antisense transcripts are retained in the nucleus while the BLV microRNAs are primordially found in the cytoplasm. How the nuclear vs. cytoplasmic localization of AS1-L and the balance between AS1-S vs. AS1-L evolves over the entire life cycle of the virus, in different cell types and in times of cellular stress remains an open question. Exploring these dynamics could be important to understand the significance of microRNA mediated cleavage of AS1-L.

Other microRNA producing viruses such as SV40 [35] and Epstein-Barr virus [36] possess viral microRNAs that cleave products from the complementary strand. It is thought that this microRNA mediated regulation manages the transition from latency (low stress) to the Lytic stage (high stress). With stress mediated suppression of microRNA production, cleavage of the antisense product is relaxed, allowing it to signal activation of the latent virus [37]. It is conceivable that such a role for the BLV AS1-S/L transcripts exists as RNA pol III transcription is known to be regulated by cell stress and cell cycle stage [38]. In addition to reducing the amount of microRNAs available for cleaving the AS1-L, changes in RNA pol III promoter occupancy and transcription could increase the fraction of AS1-L produced by removing RNA pol III complexes on the sense strand potentially impeding RNA Pol II and encouraging production of the larger transcript (AS1-L). It is apparent in Fig. 1 & Fig. S4 that antisense coverage diminishes as we enter the region containing the BLV microRNAs. RACE experiments with RNA from HeLa transfections showed that removing/inverting the microRNAs/PAS results in a higher faction of AS1-L (Fig. S8) without precipitating changes in the localization of the AS transcripts. Suggesting the removal of a insulator rather than an elimination of microRNA mediated cleavage of AS1-L and leaving the significance of microRNA mediated cleave of AS1-L unresolved.

The significant size difference between AS1-S and AS1-L is intriguing as both contain the same potential ORF. Assuming that the ORF does produce a functional protein, the role of this protein remains opaque as database searches did not reveal any obvious homology with known protein domains. In HTLV-1, a number of roles have been assigned to the HBZ protein, including the suppression of HTLV-1 5’LTR transcription and repression of the host immune response [14]. In HTLV-2 the antisense protein APH-2 also plays a role in suppressing viral transcription [39,40]. However, the potential protein in BLV AS1-S/L (87 amino acids) is considerably smaller than that seen in HTLV-1 (206 amino acids spliced / 209 amino acids unspliced) and HTLV-2 (183 amino acids) which argues against an exactly analogous protein (although the final effect may be equivalent). In addition to the roles ascribed to the HBZ protein, it has been reported that the *HBZ* mRNA is capable of supporting proliferation [18,19] and is primarily retained in the nucleus [41], suggesting a lncRNA like function. Only a small portion of AS1-L is occupied by the potential ORF and the AS1-L\S transcript is also primarily retained in the nucleus, again hinting at a potential lncRNA like role. Recent years have seen a rapid expansion in our understanding of the numerous roles lncRNAs play in the cell [42]. Many of these lncRNAs also show a predominately nuclear localization and are involved in transcriptional regulation [43]. Additionally it is also apparent that many transcripts previously described as lncRNAs possess small ORFs (<100 amino acids) that do not resemble those found in well characterized mRNAs [44]. The situation for the BLV antisense transcripts appears similar. It is thus reasonable to assume that a main role of the BLV AS1-S/L is likely to be silencing of the 5’LTR. Future work using in-vitro infection with altered proviruses will be needed to firstly establish the precise role AS1-S/L plays in the life cycle of BLV and secondarily if these different roles are carried out via the RNA or potential protein product. With this information in hand it will be interesting to compare and contrast the roles AS1-S/L and *HBZ* play and given the tractability of the BLV model, make targeted changes to the transcript and examine the effect on tumor development.

Looking beyond AS1-S/L their remains the enigma of the AS2 transcript, where at the moment the potential functions are even more obscure than for AS1-S/L. No comparable antisense transcript is seen in HTLV-1 and no potential ORF is seen in the transcript. Furthermore, the expression levels of AS2 are lower than that observed for AS1 (Fig 1 & Fig. S4). Additionally, the second exon of AS2 is small (125bp) creating a transcript containing just over ∽400bp of sequence from the BLV genome. However, the consistent splicing of AS2 with the host genome results in the production of a fusion transcript containing ∽400bp from BLV and additional sequences from the host genome. As a result the AS2 transcript size and composition will differ from cell to cell depending on provirus location in the host genome. It has recently been reported in HTLV-1 that antisense transcripts also form fusion transcripts with the host genome [15].

A striking feature of many HLTV-1 and BLV proviruses is the large deletions frequently observed in tumors, that can remove over half the proviral DNA, often including the entre 5’LTR and many vital ORFs, while always retaining the 3’LTR [15,20,45]. These observations strongly indicate that *HBZ* plays an important role in leukemogenesis and the similar pattern of deletions in BLV points to analogous role for the BLV AS1-S/L. The case of the BLV provirus T1345 is intriguing for this reason as the large deletion gives us a clue about which BLV antisense transcripts are important in the tumor. In this provirus a ∽4.3 kb deletion removes many essential viral genes but retains the BLV microRNAs [20] and continues to display robust antisense expression (Fig. 7). This deletion also removes the second exon of AS2 and the last ∽100bp of AS1-L indicating that these transcripts/regions are dispensable in the tumor.

**Figure 7.**
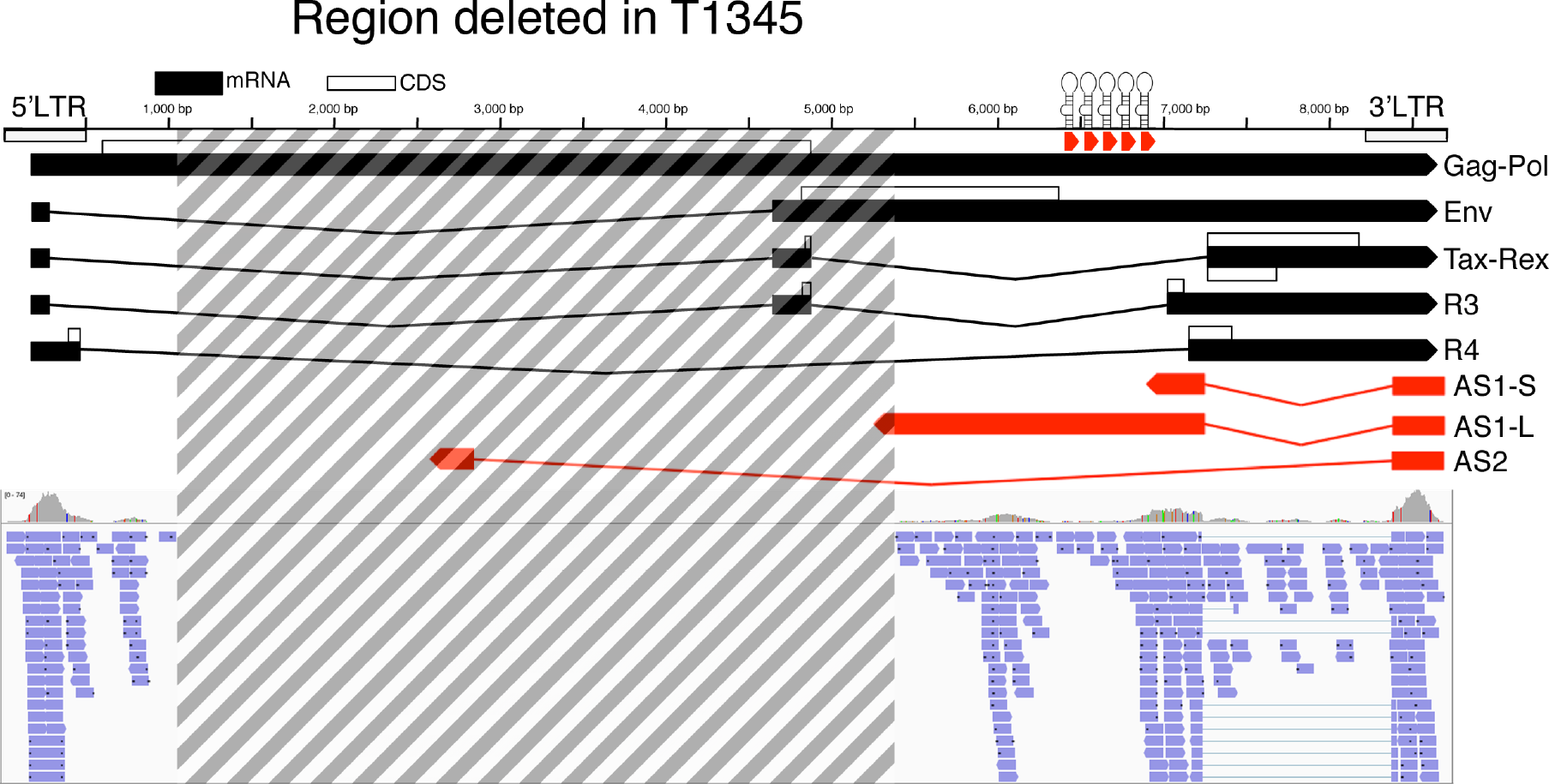
Continued antisense transcription in the defective provirus T1345. Ideogram showing the 4.4kb region deleted (dashed lines) in the T1345 provirus, identified in a monoclonal bovine tumor originating from a natural infection. Below is a screen shot from IGV showing that despite the deletion and the removal of the potential PAS for AS1-L, antisense transcription remains robust.

Knock down is generally one of the first experiments that comes to mind when exploring the function of a transcript. However, the YR2 and L267 B cell lines are refractory to transfection, making the introduction of shRNA/siRNA problematic and requiring alternative delivery systems. To overcome this hurdle we utilized LNAs as they can be delivered to cells via unassisted uptake, removing the need for transfection. Additionally as they depend on the action of RNAse H for knockdown, they are generally more effective on transcripts primarily found in the nucleus. However while unassisted uptake is straightforward, it does require much higher LNA levels (final concentration required: 1-50 nM for transfection, 1-50 μM for unassisted uptake) making optimization and long-time courses cost prohibitive. As a consequence, we carried out a limited number of assays and designed our experiment to examine if knock down of AS1-S/L could lead to the re-activation of viral transcription from the 5’LTR. Our experiments showed that at the 5μM concentration there was an apparent reduction (but not significant) in AS1-S/L levels, while at the 10μM concentration the reduction was more pronounced and reached significance. In the case of the assay for the sense transcript Tax no expression was observed in the different treatment conditions and based on final cell counts there was no apparent effect on cell viability (Fig S9.) It has been known for some time that the lack of 5’LTR based expression is imposed via histone repressive marks and DNA methylation at the 5’LTR [22,23] and we hypothesized that if the AS1-S/L has a role in maintaining this state its knock down would result in reactivation of sense transcription. However the results outlined above indicate that knock down of AS1-S/L is not sufficient to achieve reactivation in these cell lines. More prolonged and complete knock down may have produced an effect, however given the large quantities of LNAs required for unassisted uptake this was not practical. Previous work by Colin et al [22] showed that following experientially induced chromatin remodeling in the YR2 cell line there is a rapid return to a closed chromatin conformation. It may be that the AS-1S/L transcripts play a role in this process and in order to observe any effect on 5’LTR based expression, AS1-S/L knock down must be coupled with prior opening of chromatin. It will be interesting to see if future experiments can combine LNA treatment against AS1 with treatment to open chromatin and an examination of the patterns of epigenetic marks observed at the BLV 5’LTR.

In summary, the identification of the antisense transcripts taken in conjunction with the recent identification of the BLV microRNAs [20,21] represent a major shift in our understanding of the BLV induced leukemia/lymphoma. In contrast to the prevailing paradigm of a silent provirus, our work revealed that the BLV provirus is a prodigious producer of viral microRNAs and constitutively expresses antisense transcripts in all tumors examined by us. This mirrors the situation in HLTV-1 induced ATLs where the antisense transcript HBZ has been found in all ATLs examined to date [18,46]. While much work is still required to elucidate the precise roles of the BLV antisense transcripts, it is hoped that the greater tractability of the BLV model will provided key insights into deltaretrovirus induced leukemia and provide a test bed for exploring treatment options.

## Materials and Methods

### Ovine/bovine samples and cell lines

Primary ovine and bovine tumors utilized have been described previously [20,47]. Briefly, the ovine samples came from the acute stage of the disease, with tumor development occurring 15 to 48 months post infection. Animals had been infected with either the infectious clone pBLV344/pBLVX3C or with peripheral blood mononuclear cells (PBMCs) isolated from BLV-infected animals. Experiential procedures followed national and international animal care guidelines and were approved by the Comité d’Ethique Médicale de la Faculté de Médecine Université Libre de Bruxelles [20]. The bovine samples were part of a tumor collection sourced from Belgium, France, United States and Japan maintained at −80^o^C and described previously [47]. The tumors samples formed part of B-cell lymphoid masses the results of natural infection with BLV and as a consequence the time between infection and tumor development was not recorded.

The preleukemic sheep samples were obtained from animals infected with the molecular clone pBLV344 following experimental procedures approved by the University of Saskatchewan Animal Care Committee, following Canadian Council on Animal Care Guidelines (Protocol &#x0023;19940212). The animals were housed at the Vaccine and Infectious Disease Organization (VIDO-Intervac; Saskatoon, SK. Canada). Blood was collected in EDTA Vacutainers (BD, NJ, USA) and peripheral blood monocnuclear cells (PBMCs) were isolated by layering buffy coat cells on a one-step Percoll gradient [48].

The ovine B-cell tumor cell lines YR2 and L267 were derived from the M395 B-cell leukemia and the T267 B-cell lymphoma respectively. Both cell lines share the same proviral integration site as the primary tumor they were derived from and show an absence of viral messenger RNAs originating in the 5’LTR. L267LtaxSN and YR2LtaxSN are the products of transduction of the L267 and YR2 cell lines with the pLTaxSn retroviral vector, producing constitutive expression of the viral transactivator Tax and as a consequence constitutive expression of the viral mRNAs [13]. The lymphoid cell suspensions were cultured in OPTIMEM medium (Invitrogen) supplemented with 10% FCS, 1 mM sodium pyruvate, 2 mM glutamine, MEM nonessential amino acids solution 1X (Gibco) and 100 μg/mL kanamycin at a concentration of 10^6^ cells/mL. HeLa cells were procured from the American Type Culture Collection (ATCC), while the 293T cells were a gift from F. Kanshanchi (George Mason University, Manassas, VA). HeLa and 293T cells were cultured in DMEM supplemented with 10% FCS, L-glutamine and penicillin/streptomycin.

### RNA sequencing

Total RNA was either extracted using TRIzol (Life Technologies) and treated with turbo DNAse (Life Technologies) or with Qiagen AllPrep DNA/RNA/microRNA kit. Using 1ug of RNA strand specific, ribosomal RNA depleted libraries were prepared with the Illumina TruSeq Total RNA stranded kit. The resultant libraries were assessed on an Agilent Bioanalyser DNA 1000 and by qPCR with the KAPA kit (KAPA biosystems) followed by sequencing on an Illumina HiSeq 2000 (2 x 100bp paired-end reads), generating approximately 60 million raw paired-end reads per library. Small RNA libraries were prepared using the Illumina TruSeq Small RNA Library Preparation Kit and sequenced on an Illumina NextSeq 500 (75bp single reads). Custom host-provirus reference genome builds were generated using the bovine UMD3.1 [49] or ovine OAR3.1 [50] and the proviral genome sequence for BLV YR2 (NCBI Accession: KT122858) and the resultant RNA reads were mapped to the appropriate genome using STAR (v2.3.1.u) [51] and BWA [52]. Sorting, indexing and separating of the sense from the antisense was carried out with SAMtools [53] and BAMtools [54]. Read quantification and read count normalization was carried out with FeatureCounts [55] and R packages DESeq2 [56] respectively. The percentages of AS1-S vs AS1-L transcripts observed were calculated by counting the reads aligning to the AS1-L region (5237-6895) and normalized to the region length (1658bp). Reads aligning to the region in common to AS1-S & AS1-L (6895-7217) were normalized to the length (322) followed by subtraction of the reads from the AS1-L. The fraction of AS1-S compared to AS1-L was then computed via: AS1.S.norm / (AS1.S.norm+AS1.L.norm). This was computed based on all the ovine and bovine samples as well as the YR2 and L267 cell lines. The Integrative Genomic Viewer (IGV) was used for visualization of aligned sequences [57]. Reads originating from either of the two BLV LTRs aligned to both the 5’ and 3’ LTRs due to the identical sequences of both LTRs.

### End point PCR

RNA and DNA was extracted using the Qiagen AllPrep DNA/RNA/microRNA kit. Reverse transcription was carried out using SuperScript III Reverse Transcriptase (Life technologies), with subsequent RNase H treatment (New England Biolabs) and primed with random hexamers. Each PCR reaction contained 1ul cDNA, 3 pmol forward and reverse primer, 0.1ul dNTPs (10 mM), 0.075ul GoTaq Hot Start Polymerase (Promega), 2ul Q5 buffer and 6.2ul H2O. Annealing and extension conditions were based on the manufacturer's recommendations with a total of 35 cycles used. PCR products were visualized on a 1.5% agarose gel. PCR primers used are listed in Table S5.

### High-throughput sequencing of BLV integration sites

In order to ensure that preleukemic animals were actually still in the polyclonal stage of infection we carried out high throughput sequencing of proviral integration sites. The method used is similar to that outlined by [8,58], while incorporating a number of changes to increase sensitivity and facilitate sample multiplexing and is detailed in (Rosewick et al 2016 submitted). Briefly, 5 ug of DNA is sheared in a Bioruptor Pico (Diagenode) following the manufacturer's instructions for fragments of ∽1000bp. Samples are end repaired and dA tailed with the NEBNext Ultra End Repair/dA-Tailing Module (New England Biolabs) followed by ligation to a linker containing the Nextera Reverse sequence and a upstream sequence to facilitate nested PCR. Nested PCR was carried out with primer matching the linker and the BLV LTRs. Nextera XT indexes (Illumina) were added followed by sequencing on a Illumina MiSeq instrument with 2x75bp reads (Reagent Kit v3) or 2x150bp (Reagent Kit v2). The resultant paired-end reads were aligned to the host-provirus hybrid genome with BWA. Reads were trimmed based on average base quality (>= 30) and paired end reads spanning the LTR-host junction (Read 1: 8 nucleotides mapping to the LTR; Read 2: host alignment with maximum 3 mismatches) were extracted based on soft-clipped sequences and mis-paired reads. Read numbers were determined for each proviral integration site using in-house R and Perl scripts. Clone abundance was calculated as an average for the 3’ and 5’LTRs.

### Proviral load quantification

DNA was extracted with the AllPrep DNA/RNA/microRNA (Qiagen). Proviral load was quantified using PrimeTime qPCR Assays (Integrated DNA Technologies) targeting the BLV provirus and RPS9 in the host genome for normalization (Listed in Table S5). Reactions were carried out in a 10ul volume with 50ng of template DNA, 1 X Taq Man Universal PCR Master Mix, No AmpErase UNGa (ThermoScientific) and 1 X of the appropriate PrimeTime assay mix. Thermocycling conditions were 10 min at 95°C, with 40 cycles at 95°C for 15 sec and 60°C for 1 min. Standard curves were generated using serial dilutions of DNA from the YR2 cell line, Proviral load in % PBMCs = (Sample Average Quantity) x 2 / (Sample RPS9) * 100. (In the YR2 cell line the chromosome into which the BLV provirus integrated has been duplicated, as a consequence each cell carries two copies of the provirus.)

### Identification of 5’ and 3’ ends with RACE

Total RNA was extracted from the YR2 cell line with TRIzol (Life Technologies) and treated with turbo DNAse (Life Technologies). The GeneRacer Kit (Life Technologies) was used to amplify 5’ ends of both BLV antisense transcripts with a modified protocol. The manufacturer's protocol was followed until the point of the 1^st^ PCR. At this stage primers tailed with a sequence corresponding to the Nextera forward and reverse were used (adapting the protocol described for the perpetration of 16S Ribosomal RNA Gene Amplicons). The addition of the Nextera sequences to the ends of the PCR products facilitated the addition of Illumina adapters and indexes using Nextera XT primers (Illumina), subsequent pooling and high throughput sequencing. In the modified protocol the initial round of PCR was carried out using 2.5 ul of the appropriate cDNA, 1.25ul (10^M) primer tailed with the Nextera forward or reverse (listed in Table S5), 0.5ul dNTPs (10 mM), 0.25ul Q5 High-Fidelity DNA Polymerase (New England Biolabs), 5ul Q5 buffer and 14.5ul H2O. A total of 35 cycles were carried out, extension and annealing times were based on the manufacturer's recommendations. The resultant PCR product was cleaned up using the QIAquick PCR purification kit (Qiagen), Illumina sequencing adapters and indexes were added via PCR using Illumina Nextera XT primers. The reaction was cleaned up with the QIAquick PCR purification kit (Qiagen) and the resultant library concentration was determined with PicoGreen (Invitrogen). Libraries were pooled with additional Nextera based libraries followed by sequenced 2x75bp on a Illumina MiSeq instrument.

For 3’ RACE three slightly different approaches were taken. In all cases cDNA was produced from ∽1750ng of total RNA using SuperScript III Reverse Transcriptase (Life technologies), with subsequent RNase H treatment (New England Biolabs). The primer used for cDNA priming was either the GeneRacer oligo dT or an oligo dT tailed with the Nextera reverse sequence attached (see Table S5 for oligos). The first approach to 3’RACE is similar to that that outlined in Rosewick et al submitted. Briefly the 1^st^ PCR used 5ul of cDNA as template, primers matching the BLV AS exon 1 and the Nextera reverse and Q5 High-Fidelity DNA Polymerase (New England Biolabs). Annealing temperature was 66^o^C, with a 4 min extension and a total of 25 cycles. Following cleaning with 1.8X AMPure XP beads (beckmancoulter), a second semi-nested PCR was then carried out using the same primer matching the Nextera reverse and a second primer in the BLV AS exon 1, with the same cycling conditions as above. The resultant PCR product was then sheared in a Bioruptor Pico (Diagenode) following the manufacturer's instructions for fragments of ∽400bp. End repaired and dA tailing with the NEBNext Ultra End Repair/dA-Tailing Module (New England Biolabs) was followed by cleaning with AMPure XP beads (beckmancoulter) and ligation to 60 pmol annealed oligos corresponding to the Illumina Nextera forward and reverse sequences with T4 DNA ligase (Ligase New England Biolabs). The DNA was again cleaned with 0.8X AMPure XP beads and Nextera XT indexes (Illumina) were added by PCR, the indexed libraries were again cleaned with 0.8X AMPure XP beads and quantified by PicoGreen (Invitrogen). The resultant libraries were mixed with additional Nextera based libraries and sequined at either 2x75bp (Reagent Kit v3) or 2x150bp (Reagent Kit v2) on a Illumina MiSeq instrument. Reads from both approaches were mapped and visualized in the same manner outlined for the RNA sequencing data.

The second approach to 3’RACE utilized cDNA primed with the GeneRacer oligo dT. In this approach only a single round of PCR was carried out with a primer matching the GeneRacer oligo tailed with the Nextera Forward sequence and a second primer just upstream of the potential poly A site in AS1-L tailed with the Nextera Reverse sequence (see Table S5 for oligos). The PCR used 2.5ul of cDNA, Q5 High-Fidelity DNA Polymerase (New England Biolabs), with an annealing temperature of 67^o^C and a 45 sec extension and a total of 35 cycles. The reaction was cleaned up with a MiniElute column (Qiagen). Addition of the Nextera XT indexes (Illumina) quantification and sequencing on the Illumina MiSeq instrument was carried out in the same manner as described above.

The final approach to the 3’RACE utilized the MinION's (Oxford Nanopore Technologies) ability to produce long reads to observe nearly full length AS1-L transcripts. Starting with 2.5 ul of cDNA (primed with the GeneRacer oligo dT), a 1^st^ PCR was carried out using LongAmp Taq DNA Polymerase (New England Biolabs) Cycling conditions followed the manufacture recommendations with an annealing temperature of 60^o^C, an extension time of 11 min and 35 cycles. The resultant PCR product was cleaned up twice using 0.8X AMPure XP beads to ensure the removal of small fragments. A nested PCR was then carried out, again following the manufacturer's recommendations for a 50ul reaction with 1ul of the previous PCR as template. Annealing temperature was 60^o^C, with an extension time of 3 min and 15 cycles. The resultant PCR was divided, with 10ul loaded on a 1% agarose gel stained with SYBR Safe (Thermofisher) and the remaining 40 ul loaded on a second 1% agarose, both gels were also loaded with 10kb to 200bp DNA ladder. The gels were run in parallel in the same electrophoresis tank, the stained gel was visualized under UV light and used to determine when the fragments had separated sufficiently. Using this gel as a guide the portion of the unstained gel containing fragments between ∽1.5kb and 5kb was excised and purified using the MinElute Gel Extraction Kit (Qiagen). This was done to avoid exposing the DNA fragments to UV light as they will be directly sequenced on the MinION device. Using 1ug of the purified size selected DNA as template a library was prepared following the Amplicon sequencing v11 protocol, with the SQK-MAP-006 Nanopore sequencing kit followed by analysis using the SQK-MAP-006 protocol. FASTA sequences for both the 2D reads (where both strands of the fragment are sequenced) and 1D reads (only one strand sequenced) were extracted with Poretools [59]. The sequences were mapped to the same custom hybrid Ovine/BLV genome described above in the RNA sequencing section using the option for Nanopore reads in BWA [60], followed by processing with SAMtools [53] and visualization in IGV [57].

### Luciferase assays

The BLV LTR utilized was derived from the pBLV344 plasmid described by Sagata et al 1985 [61]. The various constructs were cloned into a pGL3 basic luciferase reporter plasmid. Primers used to amplify the LTR from pBLV344 and for site directed mutagens are listed in Table S5. HeLa cells were transfected in triplicate using lipofectamine 2000 with 400 ng of the appropriate construct in addition to 10ng Renilla control plasmid. In cases where the size of insert differed, amount of DNA transfected was adjusted to ensure equal copy numbers of each construct. Tax expression constructs used were previously described [12]. Forty-eight hours post transfection the cells were processed using the Dual-Glo Luciferase Assay System (Promega) following the manufacturer's instructions. Statistical significance was assessed via Tukey's multiple comparisons test, carried out using the Prism software (Graphpad), p-value < 0.05 was considered statistically significant.

### Protein coding potential and nucleotide diversity

Coding potential was assessed using the Coding-Potential Assessment Tool (CPAT) [28]. The limited number of annotated bovine lncRNA made the construction of a robust bovine specific model impractical. Instead full length sequence of the BLV antisense transcripts, five protein coding genes from HLTV-1 and BLV (including HBZ) along with 220 bovine and ovine lncRNAs obtained from rnacentral.org [62] were collected. These sequences were then analyzed with the CPAT web server (http://lilab.research.bcm.edu/cpat/) using precompiled models based on Human and mouse training sets [28].

In order to determine the sequence conservation of the potential open reading frame in AS1-S/L we examined the region in a number of BLV genomes. BLV consensus sequences were extracted from ovine and bovine aligned RNA-Seq data (STAR) using a samtools mpileup, bcftools and seqtk based custom script. An additional six BLV whole genome sequences were downloaded from NCBI (http://www.ncbi.nlm.nih.gov/nuccore) and included in the analysis. Calculation of percentage of identity and significance carried out in the same was as described in Rosewick et al [20].

### Sub-cellular location of BLV antisense transcripts

To produce cytoplasmic and nuclear fractions for YR2 we followed the method outlined in Weil et al [63]. Briefly, ∽40x10^6^ cells were washed twice in PBS, 200ul of fractionation buffer (Tris HCl 10 mM, NaCl 140 mM, MgCl2 1.5 mM, EDTA 10 mM, NP40 0.5%, RNaseOUT 100 U/ml) was then added to the cells and incubated on ice for 5 minutes. The resultant lysate was centrifuged for 5 min at 4^o^C and the supernatant transferred to a new tube (retaining the pellet), followed by centrifugation at full speed for one minute. The supernatant was again transferred to a new tube and 1 ml Trizol added (cytoplasmic fraction). For the pellet, 200ul of fractionation buffer was added and the pellet mixed by pipetting, followed by centrifugation for 5 min at 4^o^C. The supernatant was discarded and the pellet resuspended in 200ul of fractionation buffer followed by disruption in a TissueLyser II (Qiagen), with the subsequent addition of 1 ml of Trizol (nuclear fraction). RNA was extracted from both fractions and the resultant RNA was then treated with turbo DNAse (Life Technologies). RNA from both fractions was sequenced and the reads mapped in the same manner as outlined above. Enrichment in nuclear and cytoplasmic fractions was computed as follows : Paired-end RNA-Seq reads were aligned on the human (hg19) or ovine (OAR3.1) with STAR [51]. Read counts for the different RNA species (snRNA,snoRNA,tRNA,.) were computed using featureCounts and ENSEMBL v74 annotation. In order to take into account sequencing depth variation across samples, read count normalization was performed using DESeq2 [56]. Finally, abundance of each RNA species was computed by comparing the total normalized read count of each RNA species.

Real time PCR based absolute quantification (see Table S5 for oligos) was carried out on the ABI Prism 7900HT Sequence Detector System (Applied Biosystems) using ABsolute Blue QPCR SYBR Green ROX Mix (Thermo Scientific). PCR products were cloned into a TOPO TA plasmid (Life Technologies) and then diluted to the appropriate concentration to produce a standard curve. Relative enrichment of the transcript was calculated in the same manner as Kobayashi-Ishihara et al [64] % enrichment = RNA level in nucleus / combined RNA levels for nuclear and cytoplasmic fraction x 100.

### Identifying products of RISC mediated cleavage

RNA was extracted from the YR2 cell line and a preleukemic sheep (17 months post inoculation with pBLV344) using Trizol (Life Technologies) and treated with turbo DNAse (Life Technologies). In the case of the cytoplasmic and nuclear RNA YR2 cells were fractionated following the method outlined above. The procedure for identifying the products of slicing was based on Davis et al [32]. The RNA Oligo from the gene RACE kit (Promega) was ligated to 3.8ug of total RNA using 1ul 10x Ligase Buffer, 1ul 10mM ATP, 1ul RNaseOut, 1ul T4 RNA ligase in a total volume of 10ul. The mix was incubated for 1hr at 37^o^C followed by precipitation and resuspension of the RNA in 10ul of H2O. Revere transcription was carried out with Revert Aid Premium Reverse Transcriptase (Thermo scientific) and random hexamers. PCR was initially carried out in five separate reactions with the common forward Gene Racer 5’ primer in combination with a primer designed to identify the slicing of one of the BLV microRNAs. The PCR reaction contained 2ul cDNA, 0.3ul (10 ¿vM) forward and reverse primer, 0.1ul dNTPs (10 mM), 0.1ul Q5 High-Fidelity DNA Polymerase (NEB), 2ul Q5 buffer and 5.2ul H2O (see Table S5 for oligos). A total of 30 cycles were performed using annealing and extension conditions recommended by the manufacturer. A nested PCR was then carried out using tailed primers carrying the Nextera forward and reverse sequences. The resultant PCR products were pooled and cleaned up with the QIAquick PCR purification kit (Qiagen). As outlined above the Illumina adapters and indexes were added using Nextera XT indexes (Illumina). The resultant libraries were paired-end sequenced 2 x 75bp on an Illumina MiSeq using the Reagent Kit v3. To determine the precise position of the free 5’ end of the RNA we took the 30bp sequence of the RNA oligo and concatenated it to a sliding window of 31bp sequences from the BLV microRNA region. Starting at position 6357 a total of 554 61bp hybrid sequences were created. A BASH script was then used to search the FASTA files of each library for perfect matches with the concatenated sequences, thereby identifying the position and frequency of RNA cleavage. The high throughput sequencing reads were also mapped to the ovine-BLV hybrid genome using BWA and visualized in IGV.

### Deleting and inverting the BLV microRNAs

The removal and inversion of BLV microRNAs was carried out on the wild type molecular clone pBLV344 [65]. PCR primers were designed to amplify a 3kb region just upstream of the BLV microRNA cluster and a 1.7 kb region downstream of the microRNAs that included the entire terminal portion of the BLV provirus. The recognition sequence for the MluI restriction enzyme was used to tail the reverse primer of the upstream primer pair and the forward primer of the downstream pair (see Table S5). PCR was carried out using Phusion High-Fidelity DNA Polymerase (NEB) with the pBLV344 plasmid as template. PCR products were digested with MluI (NEB) and then ligated together using promega T4 DNA ligase. The ligation product was run on a 1% gel and the band of the appropriate size (4.7kb) cut out and purified. The altered BLV fragment was then introduced into the pBLV344 plasmid using the BspEI (NEB) and NheI (Promega) restriction enzymes. This resulted in a provirus lacking the BLV microRNAs with a MluI recognition site incorporated in their place. The inverted plasmid was created by amplifying the BLV microRNA cluster using primers tailed with the MluI recognition site. The resultant PCR product and the pBLV344 plasmid lacking the microRNAs (with MluI site added) were digested with MluI and the PCR product inserted into the provirus by ligation. Sanger sequencing was used to check the orientation of the reinserted region and indentify a plasmid where the BLV microRNA orientation was inverted. The resultant constructs were transfected into HeLa cells followed by RNA extraction, using the same protocol outlined above.

### Antisense knock down with locked nucleic acid antisense oligos (LNAs)

The BLV cell lines YR2 and L267 are refractory to transfection, as a consequence it was not possible to knock down the expression of the BLV antisense transcripts using more conventional approaches such as siRNAs. We therefore employed locked nucleic acid antisense oligos (Exiqon), introduced via unassisted uptake [33], following the manufacturer's recommendations. 500ul of YR2 and L267 cells at 10^6^ cells/mL were placed in 24 well plates. Each cell line was treated in duplicate with a mix of three LNAs, that targeted different parts of AS1 (Table S5). Two different final concentrations were tested, 5 |iM and 10 |iM. As a negative control both cell lines were treated in duplicate with the LNA longRNA GapmeR Negative control A (Exiqon), at a final concentration of 10 | M, a mock treatment using 25 ul of H20 was also carried out. The cells were incubated for 5 days and cell numbers were then estimated with a Bio-Rad TC20™ Automated Cell Counter. DNA and RNA were extracted using the Qiagen AllPrep DNA/RNA/microRNA kit. cDNA was produced using SuperScript III (Invitrogen) and random hexamers. Real time PCR was carried out using PrimeTime qPCR Assays (Integrated DNA Technologies) and Taq Man Universal PCR Master Mix, No AmpErase UNGa (ThermoScientific) on a ABI 7900HT Fast Real-Time PCR System. Data analysis was carried out with the qbase+ software (Biogazelle), with graphs and statistical analysis carried out using the Prism software (Graphpad). Expression levels for each sample were normalized against RSP9 and TBP levels. In order to combine the expression levels for both cell lines, expression levels were first scaled to the LNA Negative control of each cell line. P values of ≤ 0.05 were considered to be statistically significant after controlling for multiple testing. Statistical significance was assessed via a Kruskal-Wallis test and Dunn's multiple comparisons test, P values of ≤ 0.05 were considered to be statistically significant.

## Acknowledgements

This work was supported by the Fonds de la Recherche Scientifique - FNRS under Grant n° 3.4587.12, Télévie, les Amis de l’Institut Bordet, the International Brachet Stiftung (IBS), the Fondation Lambeau-Marteaux, and a Télévie Grant to VH. KD is Postdoctoral Researcher of the FNRS, NR and MA are Scientific Research Worker of Télévie. We thank the GIGA Genomics Platform and more particularly Wouter Coppieters, Benoit Hennuy and Latifa Karim for their help with high-throughput sequencing.

